# Purified diets containing high levels of soluble fiber and grain-based diets promote similar gastrointestinal morphometry yet distinct microbial communities

**DOI:** 10.1101/2024.04.08.588600

**Authors:** Elaine M. Glenny, Jintong Liu, Harlyn G. Skinner, Tori L. McFarlane, Kylie K. Reed, Alyssa Weninger, Zorka Djukic, Michael A. Pellizzon, Ian M. Carroll

## Abstract

Grain-based diets (GBDs) are widely used in rodent studies but their utility is limited due to batch-to-batch variability resulting from inconsistent ingredients. Purified diets (PDs) are composed of only known and refined ingredients and offer a solution to the constraints of GBDs. However, unlike GBDs, PDs commonly used as control diets typically contain little to no soluble fiber. We therefore sought to identify a combination of fibers in PDs that best recapitulates the gastrointestinal morphometry and intestinal microbial composition of mice fed GBDs. Adult male mice (n=30) were randomly assigned to one of six diets—two GBDs and four PDs with varying insoluble and soluble fiber composition—for 28 days. 16S rRNA gene sequencing was used to compare microbial profiles across different gastrointestinal (GI) niches and diets. Gut microbiotas and cecal weights were distinct between mice fed the two GBDs, indicating that GBDs are unreliable controls in diet-based studies. Unexpectedly, intestinal microbial richness decreased as the amount of soluble fiber in the PDs increased and the addition of multiple soluble fibers did not rescue this effect. Mice fed PDs with high soluble fiber content (≥ 75% of dietary fiber was soluble fiber) best recapitulated GI morphometry of mice fed GBDs, but intestinal microbial communities were distinct between PD- and GBD-fed mice. Although supplementing PDs with soluble fiber improved GI morphometry, further research to determine the optimal mixture of soluble and insoluble fibers is required to more closely mirror the intestinal microbial diversity observed in mice fed GBDs.

**Importance:** Dietary fibers are essential for maintaining gut health. Insoluble fibers aid in fecal bulking and water retention while soluble fiber is a fermentative substrate for intestinal microbial communities. GBDs are commonly used in preclinical research but the variability in ingredients across batches impedes reproducibility. PDs, which are composed of highly refined ingredients, pose a potential solution but the most widely used low-fat control PDs contain no soluble fiber. This study intended to identify a PD with a combination of fibers that promotes murine gut health and microbial diversity. A PD with optimal fiber composition would aid in the standardization and reproducibility of studies investigating intestinal physiology and the gut microbiota.

## INTRODUCTION

Intestinal microbial communities influence gastrointestinal (GI) physiology in health and disease (1–3). Multiple host and environmental factors including age, geographic location, sex, antibiotic use, and diet impact intestinal microbial composition and consequent community functions (4). Acute diet switches and chronic dietary patterns particularly impact the gut microbiota, as even short-term dietary changes alter gut microbial communities in humans (5, 6). Additionally, preclinical research demonstrates analogous findings—dietary factors such as fat content can alter gut microbiota composition in mice (7).

Fiber is a principal dietary constituent that significantly influences intestinal microbial community structure and is critical for promoting GI health (8). Insoluble fiber is a fecal bulking and water retention agent that influences GI transit time (9). Conversely, gut microbes that encode the required enzymes can ferment soluble fibers to short-chain fatty acids to provide an energy source for intestinal microbial communities and the colonic epithelium (10). A deficit in dietary soluble fiber may necessitate enteric microbes to extract energy from the colonic mucus barrier, thereby increasing the host’s susceptibility to pathogenic infection (11, 12). Thus, fiber is a crucial nutrient for both intestinal microbial communities and the host.

Diets used in preclinical experiments are typically either grain-based diets (GBDs) or purified diets (PDs). GBDs, informally known as chow, are generally manufactured from agricultural and animal by-products such as ground corn, ground oats, alfalfa meal, soybean meal, and ground wheat (13). While economical, GBD ingredient composition—including dietary fibers—will vary across batches (13). In contrast to GBDs, PDs are defined diets with highly refined ingredients, and therefore contain precise amounts of each ingredient and nutrient to negate variability between batches and increase reproducibility across animal studies (14, 15). However, the most popular PDs, including the AIN-76A and AIN-93 series (AIN-93G and AIN-93M), contain only 5% total fiber as cellulose, which is an insoluble fiber and mostly non-fermentable (16–18). Chassaing et al. demonstrated that PDs containing only cellulose increased adipose stores and promoted abnormal GI morphometry in mice relative to mice fed a GBD, and that these observations could be reversed with the replacement (or addition) of inulin to the PDs (19)—others have noted similar findings (20–22).

The goal of the current study was to identify a combination of dietary fibers in a PD that promotes optimal murine gut morphometry and an ecologically diverse intestinal microbial community. We evaluated the benefit of different fiber mixtures by: (i) characterizing the impact of six different murine diets (two distinct GBDs and four PDs with varying fiber composition) on murine adiposity and GI morphometry; (ii) comparing the microbial composition of mice consuming these six diets across three distinct GI niches (cecum, colon, and feces); and (iii) identifying specific bacterial genera that were differentially abundant across mice fed the different diets. As balanced diets for humans contain a mix of soluble and insoluble fibers, we hypothesized that diets with a greater diversity of soluble and insoluble fibers would support the most diverse intestinal microbial community and promote normal GI morphometry in mice.

## RESULTS

### Amount and variety of dietary fiber impacts food consumption, GI morphometry, and adiposity

Cumulative food intake varied across diet groups (**Figure 1A**). Mice consuming LabDiet 5001 had a higher average daily calorie intake than mice assigned to Teklad 2020X and every other PD except 75C/25I (**Figure 1B**). Mice that consumed a greater quantity and mix of soluble fibers (25C/I/G/P) ate fewer calories relative to mice fed a greater amount of insoluble fiber (75C/25I) (**Figure 1B**).

**Figure 1.**
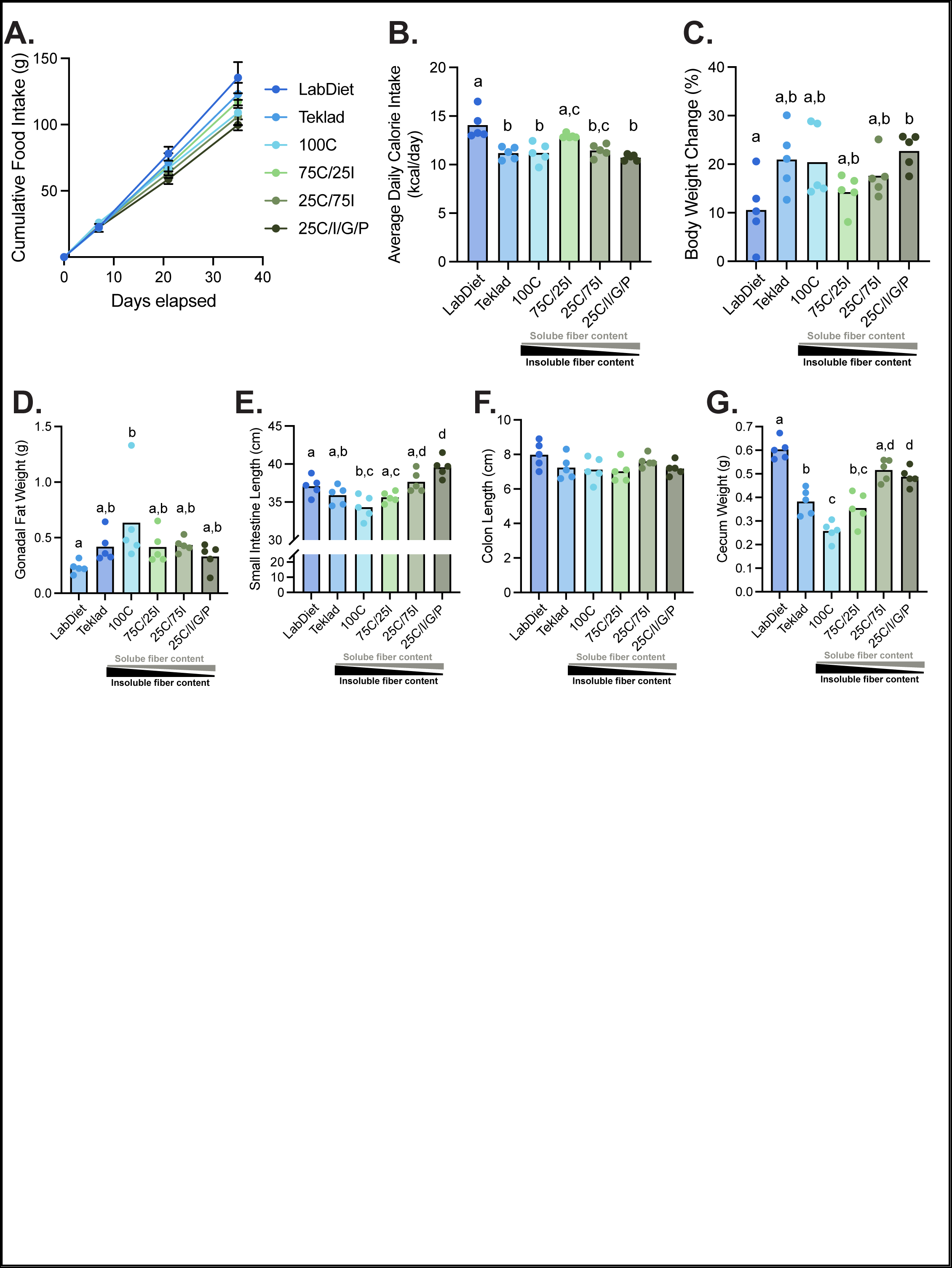
The type of amount of dietary fiber impacts food consumption, intestinal morphometry, and adiposity. Cumulative food intake **(A)**, average daily calorie intake **(B)**, percent body weight change **(C)**, gonadal fat weight **(D)**, small intestine length **(E)**, colon length **(F)**, and cecum weight **(G)** are shown for mice in each dietary group. Statistical differences were determined using a one-way ANOVA with Tukey’s post-hoc test. Different letters denote significant difference (p<0.05) between groups.

Mice assigned to LabDiet 5001 gained less weight compared with mice fed the 25C/I/G/P diet (**Figure 1C**), although no significant differences in absolute body weights was detected between diet groups at the end of the experiment. Additionally, mice assigned to the 100C diet had more gonadal fat than mice fed LabDiet 5001 diet (**Figure 1D**). No other differences were observed in body weight or adipose stores across the other diet groups.

Relative to mice fed LabDiet 5001, small intestines were longer in mice fed the 25C/I/G/P diet and shorter in those consuming the 100C diet (**Figure 1E**). Increasing the amount and variety of dietary soluble fiber impacted small intestinal length as mice fed the 25C/I/G/P diet exhibited longer small intestines than mice consuming the 75C/25I and 100C diets (**Figure 1E**). No differences were observed in colon length (**Figure 1F**). Ceca from mice fed LabDiet 5001 were heavier than mice fed Teklad 2020X (**Figure 1G**). In the PDs, increasing the amount of soluble fiber in the diet was associated with heavier ceca (**Figure 1G**).

### Increasing amounts of soluble dietary fiber reduce gut microbial diversity in a stepwise manner

Microbial communities were characterized across three GI niches at the end of the study (day 35)— cecum, colon, and feces. Microbial richness (determined by Shannon index and observed number of SV) did not differ between cecal and colonic samples; however, fecal samples exhibited greater diversity based on Shannon index (**Figure 2A-B**). Contrary to our hypothesis, increasing the amount of soluble fiber was associated with a reduction, rather than an increase, in α-diversity. In the cecum and colon, the greatest microbial diversity (based on Shannon index) was observed in mice fed either a GBD or the 100C diet lacking soluble fiber (**Figure 2C-F**). Shannon Index highlights this observation as mice ingesting the most soluble fiber (25C/75I and 25C/I/G/P) had the lowest cecal α-diversity (**Figure 2E**). Additionally, cecal samples from mice consuming Teklad 2020X had lower observed SVs compared to LabDiet 5001 (Figure 2F). In fecal samples, while there were no differences for Shannon diversity across any diet, mice fed the Teklad GBD had a greater number of observed SVs relative to mice consuming the 25C/75I PD diet (**Figure 2G-H**).

**Figure 2.**
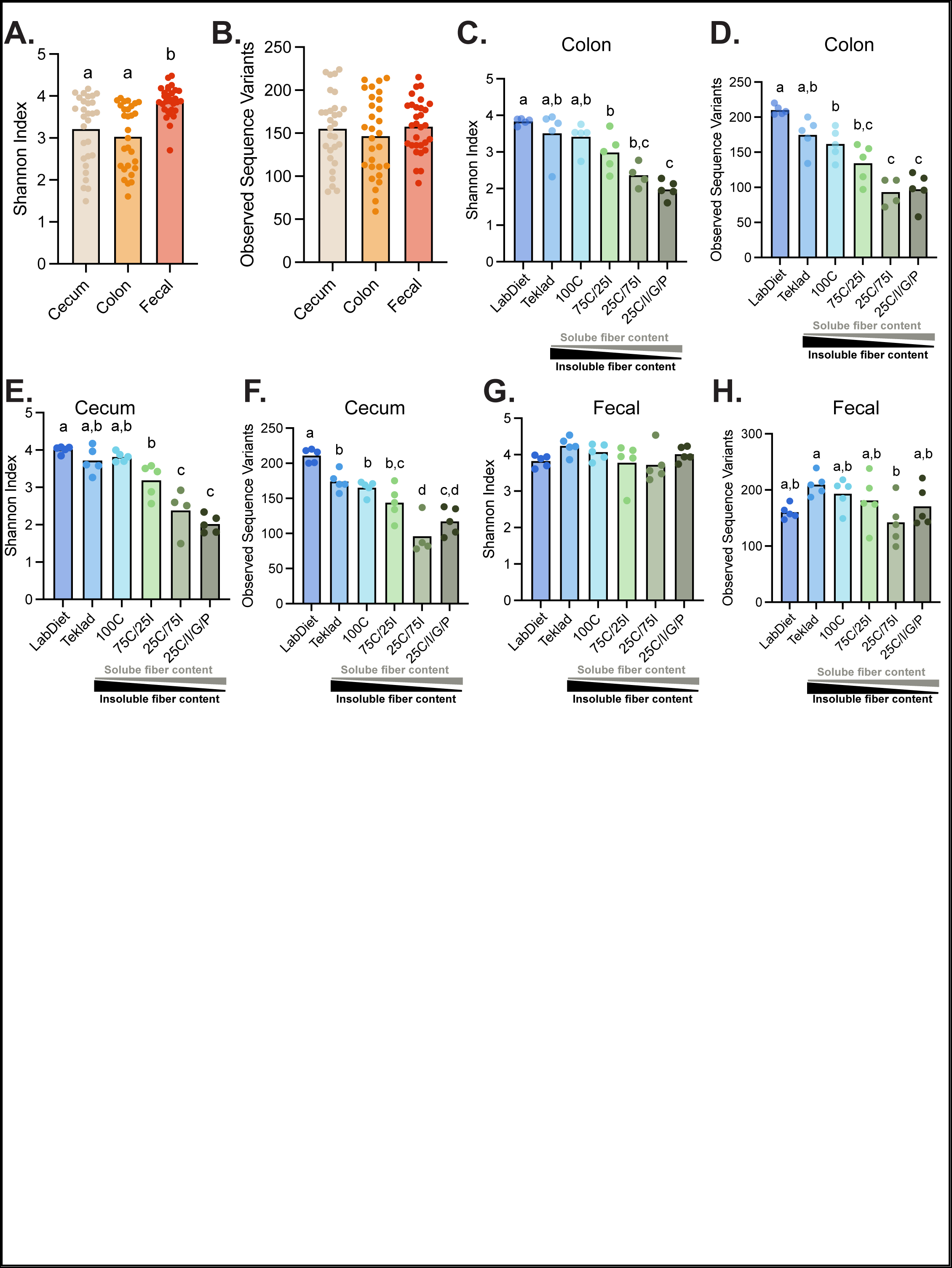
Increasing amounts of soluble dietary fiber reduce gut microbial diversity. Shannon index **(A)** and number of SV **(B)** across three GI niches. Shannon index **(C)** and number of SV **(D)** in colon samples amongst diets groups. Shannon index **(E)** and number of SV **(F)** of cecal samples amongst diets groups. Shannon index **(G)** and number of SV **(H)** of fecal samples amongst diet groups. Statistical differences were determined using a one-way ANOVA with Tukey’s post-hoc test. Different letters denote significant difference (p<0.05) between groups.

### Fiber composition shapes gut microbial communities

Multidimensional scaling (MDS) plots using Bray Curtis dissimilarity distances revealed significantly different microbial community structures in the cecum and colon compared with feces (**Figure 3A**). MDS plots combining samples across all three GI niches revealed that microbial communities were distinct between mice fed GBDs and PDs and this observation was more pronounced within the cecum and colon (**Figure 3B-E**). Mice consuming the highest amounts of soluble fiber (25C/75I and 25C/I/G/P) had different cecal and colonic microbial communities relative to mice fed PDs with less soluble fiber (**Figure 3C&D**). Adjusting the amount of soluble and insoluble fiber (75C/25I vs. 25C/75I) did not result in distinct microbial communities while reducing insoluble fiber and adding a more diverse profile of soluble fibers (25C/IG/P vs. 75C/25I) influenced microbial composition (**Figure 3C&D**). Interestingly, fecal microbial communities did not segregate based on diet group (**Figure 3E**). Overall, these data illustrate that diet type (i.e. PD vs. GBD) along with changes to soluble fiber contents (particularly increasing the number of soluble fibers) exerted a strong influence on shaping the microbial communities in the cecal and colonic niches.

**Figure 3.**
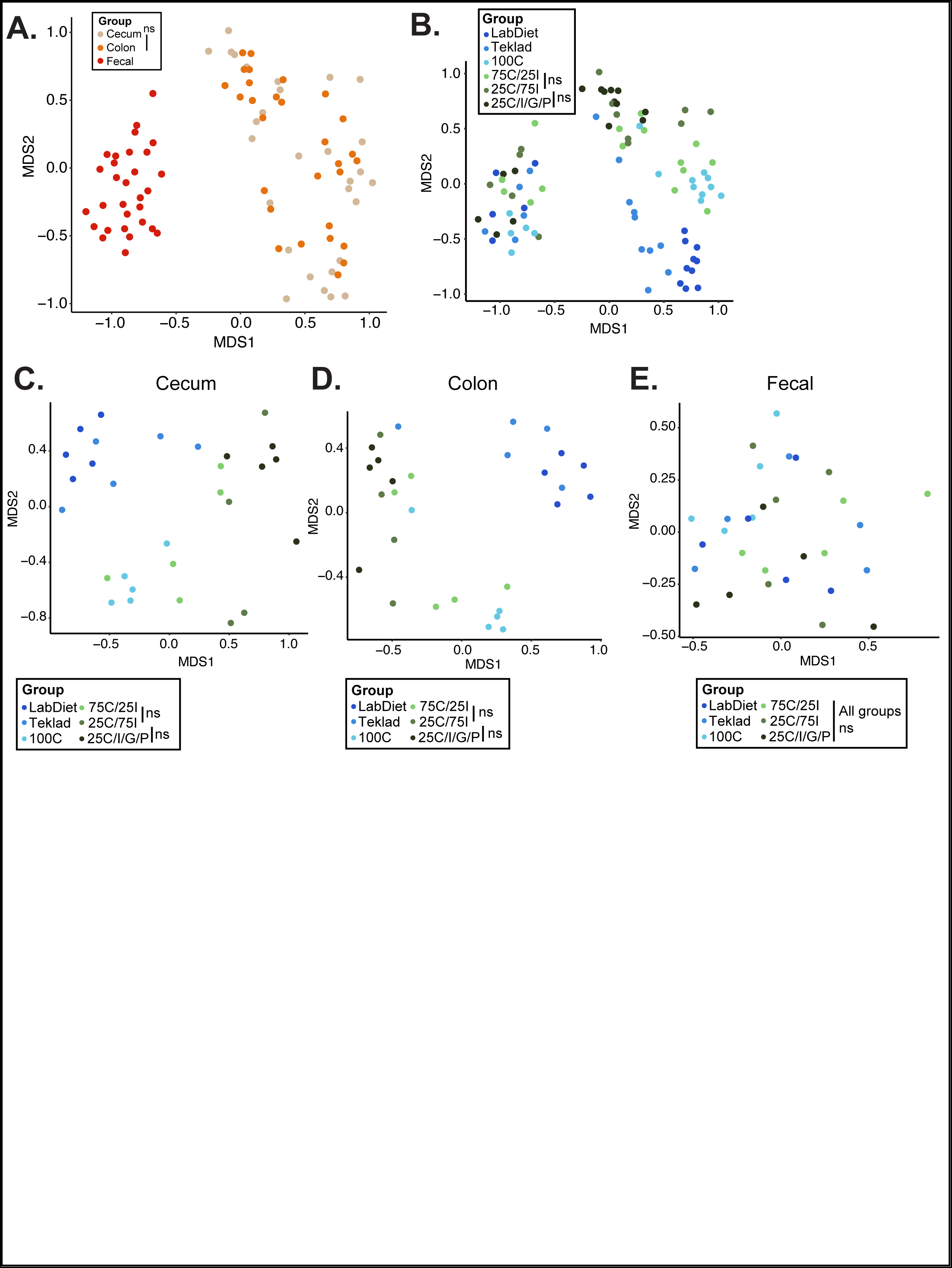
GI niche and dietary fiber influence microbial community composition. MDS plot using Bray-Curtis dissimilarity distances of microbial communities from the cecum, colon, and feces colored by diet group **(A)**. MDS plot using Bray-Curtis dissimilarity distances of all GI niches combined **(B)**, only cecal **(C)**, only colonic **(D)**, and only fecal **(E)** microbial communities colored by diet group. p<0.05 between all diet groups (PERMANOVA) unless indicated as non-significant (ns).

### GBDs differentially influence the composition of the gut microbiota relative to PDs

As cecal and colonic microbiotas exhibited a more similar composition than fecal microbial communities, and as the cecum has the greatest microbial metabolic activity, we performed a differential abundance analysis on cecal samples to identify specific bacterial genera driving differences in community composition amongst diet groups. Although the relative abundance of bacterial taxa varied between GI niches (**Figure 4 A-C**), the amount and type of fiber influenced the relative abundance of specific microbial genera (**Supplemental Table 1&2**). Based on a LEfSe differential abundance analysis, *Family XIII UCG 001*, *Lactococcus*, *Tyzzerella*, *Alistipes*, *Harryflintia*, and *Akkermansia* were the top 6 genera most likely to explain differences between groups based on the false discovery rate (FDR) q value (**Figure 4D-I; Supplemental Table 1&2**). Specifically, the relative abundance of *Family XIII UCG 001* and *Lactococcus* was higher in the cecum of mice fed PDs than mice fed GBDs (**Figure 4D-E**). The highest levels of *Tyzzerella* were found in the ceca of mice fed LabDiet 5001 (**Figure 4F**). *Alistipes* was more abundant in mice fed LabDiet 5001 relative to all other diets while *Akkermansia* was more abundant in mice fed PDs compared with mice consuming LabDiet 5001 (**Figure 4G&I**).

**Figure 4.**
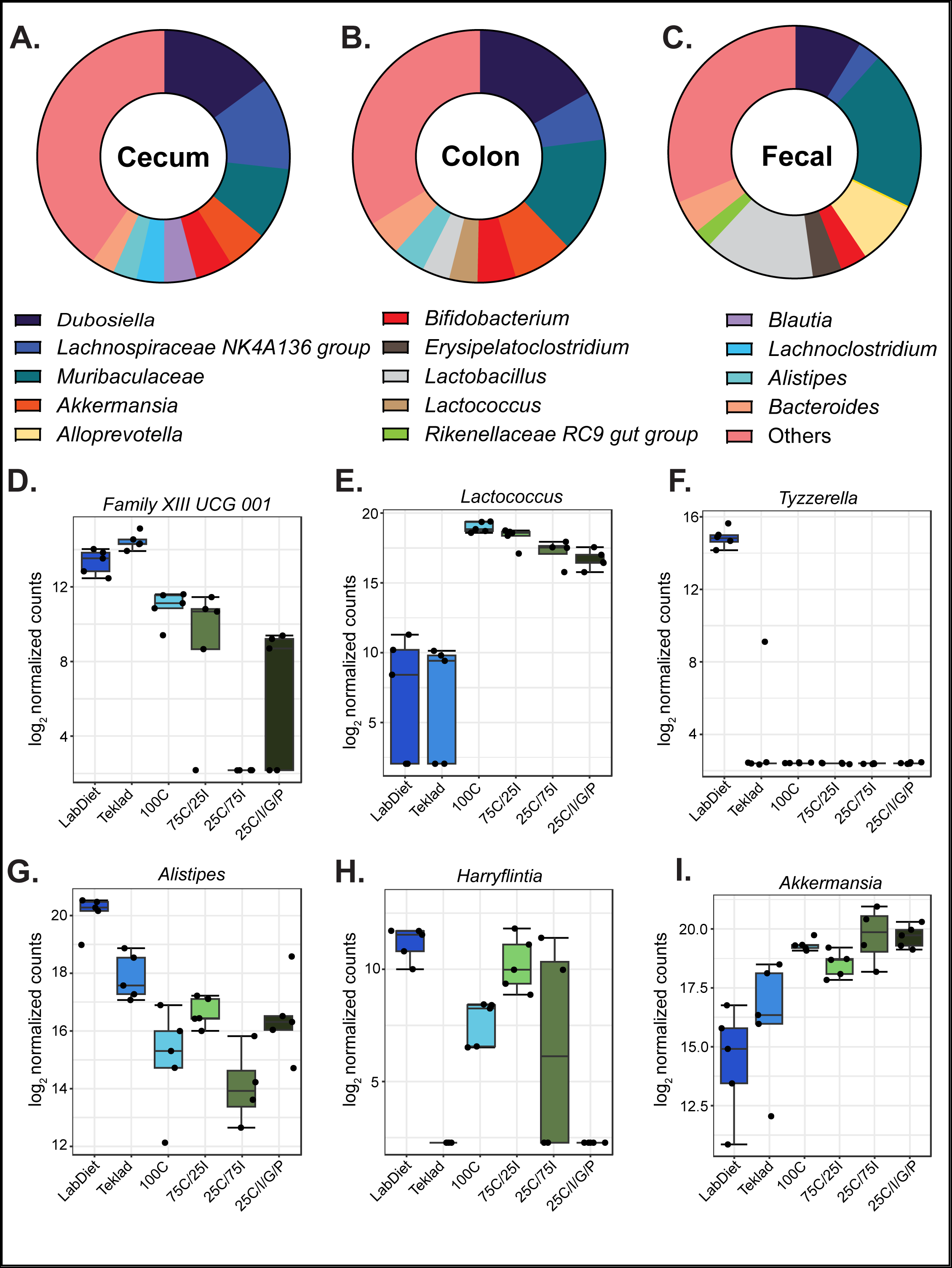
Diet types enrich for specific bacterial genera. Relative abundance of the top 9 most abundant bacterial genera in cecal **(A)**, colonic **(B)**, and fecal **(C)** microbial communities. Relative abundance in the cecum amongst diet groups for *Family XIII UCG 001* **(D)**, *Lactococcus* **(E)**, *Tyzzerella* **(F)**, *Alistipes* **(G)**, *Harryflintia* **(H)**, and *Akkermansia* **(I)**.

## DISCUSSION

Consistent with Chassaing et al., we found that mice consuming a PD lacking soluble fiber (100C) accumulated more adiposity relative to mice fed LabDiet 5001—a GBD (19). Additionally, despite consuming more calories, mice on LabDiet 5001 gained the least weight over the 28-day experimental period. Moreover, mice consuming LabDiet 5001 exhibited longer small intestines than mice fed the 25C/I/G/P diet and shorter than those fed the 100C diet. Not only did our current study reproduce major findings from Chassaing *et al*., but we also expanded upon this study by evaluating a second GBD (Teklad 2020X) to delineate whether the previous observations were specific to LabDiet 5001 or generalizable to the broader category of GBDs. Unexpectedly, unlike mice on LabDiet 5001, mice that consumed Teklad 2020X did not have less gonadal fat than mice fed any of the four PDs. We also found that although gut microbial communities clustered according to GI niche, cecal and colon microbiotas from mice consuming GBDs clustered separately from mice consuming PDs—an observation not recapitulated in the fecal microbiotas.

These results suggest diet is an experimental variable that should be carefully considered in preclinical studies. While both GBDs and PDs significantly, and uniquely, impacted weight gain, adiposity accumulation, GI morphometry, and intestinal microbial communities, open-source PDs offer the opportunity to reduce experimental variability and increase reproducibility across studies by using defined ingredients. Indeed, many rodent studies employ PDs to promote metabolic disease such as obesity, nonalcoholic steatohepatitis, and cardiovascular disease yet use GBDs for the control group (15, 23, 24). However, differences in the dietary formulations between PD and GBD— known and unknown—confound data interpretation in these studies. A recent study demonstrated that the lack of soluble fiber in PDs, rather than the addition of fat, drove alterations in the gut microbiota of mice consuming a diet commonly used to study obesity (24). Another study reported that the microbial communities between mice fed GBDs and PDs were more distinct than those between mice fed a high-fat PD and a low-fat PD (25). Our findings demonstrate that diet type (GBD or PD) significantly influences microbial community composition, with GBDs and PDs showing greater similarity within each diet type than to the other diet type. This underscores the importance of controlling for diet in these studies.

We found the addition of soluble fibers in PDs reduced microbial α-diversity in cecal and colonic microbial communities. As this observation agrees with reports demonstrating a progressive reduction in α-diversity as the contribution of soluble fiber to a diet increases, it was not surprising to find that a diet containing three sources of soluble fibers also reduced microbial diversity (21, 22). Reduced microbial diversity is typically associated with a low-fiber, high-sugar diet and linked to metabolic syndrome and inflammatory bowel diseases in preclinical and clinical studies (26). Additionally, a meta-analysis of nutritional interventions modulating dietary fibers across 64 human trials reported no significant association between fiber and microbial α-diversity (27). It was therefore not surprising that the addition of soluble fibers to rodent diets had a beneficial impact on the GI tract (based on gross morphometry) yet reduced microbial richness. One hypothesis may be that insoluble fibers, to a greater extent than soluble fibers, influence microbial richness as the PDs used in this study contained increasing quantities of insoluble fiber as the soluble fiber quantity decreased. In one report, the addition of soluble fiber sources (i.e. inulin, psyllium, or combination of both) to PDs containing cellulose had no effect on cecal microbial α-diversity in male mice following 10 weeks on diet (28). Future studies are needed to test this hypothesis and to uncover a potential mechanism.

Our study demonstrated that GI niche (cecum, colon, and feces) drove differences in microbiota composition. Additionally, gut microbiotas segregated within the cecum and colon based on whether mice were consuming GBDs or PDs. Altering the amount of soluble fiber did not have a significant impact of gut microbial community composition. Specifically, mice consuming PDs with variable amounts of soluble fiber (75C/25l, 25C/75l, and 25C/I/G/P) had distinct gut microbial communities relative to mice fed GBDs or a PD with only insoluble fiber (100C). This phenomenon was observed in cecal and colonic microbial communities, but not in the feces. The lack of diet-driven differences in gut microbiota composition within fecal samples questions the utility of this niche when investigating diet-microbiota interactions; however, the benefit of using fecal samples to investigate longitudinal changes in microbiotas may outweigh this limitation. One limitation of our study is that mucosal-associated microbial communities were not investigated. Given the proximity of mucosal-associated microbial communities to the host gut epithelial barrier (29, 30), this GI niche may better reflect the influence of intestinal microbes on the host and disease.

The *Lactococcus* genus was more abundant in the cecal contents of mice fed PDs than in mice fed GBDs. Reports have demonstrated that *Lactococcus* in rodent gut microbiotas results from the industrial production of casein, a main protein source in PDs but not in GBDs, and that the DNA is an artifact of casein production rather than detection of viable bacteria living in the intestinal tract (31). As high-throughput sequencing of the 16S rRNA gene is unable to distinguish between live and dead microbes, we are unable to determine the source of *Lactococcus* in our study but DNA fragments resulting from casein production is a potential explanation for the observed increased abundance in the ceca of mice consuming PDs relative to GBDs.

*Akkermansia* spp. are mucin-degrading bacteria that reside in the mucosal layer and exhibit anti-inflammatory properties. When broken down, mucin constitutes energy sources that can be used by the gut microbiota (32). Increased relative abundance of *Akkermansia* may reflect increased mucin production in the GI tract. As *Akkermansia* had a higher relative abundance in PDs than in GBDs, our study suggests that the PDs may promote an anti-inflammatory environment in the gut. In contrast, *Alistipes* were more abundant in mice consuming GBDs. As this genus has been reported to be beneficial and detrimental to host health (33), understanding the impact of GBDs on the gut microbiota requires further investigation.

## CONCLUSION

In conclusion, mice fed PDs with high soluble fiber content (≥ 75% of total fiber, ∼7 gm% total) best recapitulated the GI morphometry of mice fed GBDs, but no PD recapitulated the gut microbial composition of mice fed GBDs. Our results confirm that diets are an important experimental variable and GBDs are an inappropriate control for diet-based studies. Despite the limitations of using GBDs in diet-based research, our study highlights that GBDs confer benefits to GI health and microbial richness relative to PDs, even with multiple fiber sources. Further work to formulate a PD with a mixture of soluble and insoluble fibers that recapitulates these attributes of GBDs is highly desirable to improve both the reproducibility and the broader impact of preclinical rodent studies.

## MATERIALS AND METHODS

### Animals

Six-week-old male C57BL/6J mice were purchased from Jackson Laboratory (Bar Harbor, ME; n=30 mice). Mice were individually housed in ventilated cages with *ad libitum* access to autoclaved water and irradiated food. All autoclaved cages contained bedding, a hut, and a nestlet. During the seven-day acclimation period, all mice were fed the GBD Teklad 2020X. Mice were then randomly assigned to one of the six diets for 28 days (n=5 mice/group). Food consumption and body weight were measured on day 0 (when all mice were placed on Teklad 2020X), day 7 (when mice were randomly assigned to one of six diets), day 21, and day 35 (when the study terminated). Food was replaced and cages were changed on days 7 and 21. All animal care and experimental protocols were approved by the Institutional Animal Care and Use Committee (IACUC) of University of North Carolina-Chapel Hill.

### Diets

All six diets used in this study contained sufficient essential vitamins and minerals and all had a similar macronutrient profile. Descriptions of the diets, including dietary fiber compositions and caloric content, are listed in **Table 1**. In brief, the two GBDs were Teklad 2020X (Inotiv; Madison, WI) and LabDiet 5001 (LabDiet; St. Louis, MO). Based on chemical composition data from the manufacturers, Teklad 2020X, a phytoestrogen-free diet (i.e., excludes alfalfa and soybean meal) contains approximately 24 kcal% protein, 16 kcal% fat, and 60 kcal% carbohydrate; LabDiet 5001 contains both alfalfa and soybean meal and approximately 29 kcal% protein, 14 kcal% fat, and 57 kcal% carbohydrate. Fiber contents of the GBDs were analyzed by Medallion Labs (Minneapolis, MN) using the Association of Official Analytical Chemists method 991.43 (modified) to determine total, insoluble, and soluble fiber (LabDiet 5001: 21.1% total, 15.4% insoluble, 5.7% soluble; 2020X: 16.7% total, 12.3% insoluble, 4.4% soluble) (34). The four PDs were gifted from Research Diets, Inc. (New Brunswick, NJ) and were based on the Open Standard Diet D11112201 (22). The PDs contained varying compositions of insoluble fiber (**100C**: 100% of dietary fiber was derived from the insoluble fiber cellulose) and soluble fibers (**75C/25I**: 75% of dietary fiber was derived from cellulose and the remaining 25% from the soluble fiber inulin [Chicory root, Orafti HP, Beneo]; **25C/75I**: 25% of dietary fiber was derived from cellulose and 75% from inulin; **25C/I/G/P**: 25% of dietary fiber was derived from cellulose and 75% from a mix of inulin, glucomannan [Konjac root, NOW Foods], and pectin [Citrus Peel, 1400, Tic Gums], each of which contain mainly soluble fiber based on analysis by Medallion Labs). To achieve isocaloric PDs, corn starch was added as needed to replace calories derived from soluble fibers.

**Table 1.** Composition of experimental diets.

| Product # | D11112225 |  | D11112201 |  | D11112231 |  | D11112232 |  |
| --- | --- | --- | --- | --- | --- | --- | --- | --- |
| Diet Group Name | 100C |  | 75C/25I |  | 25C/75I |  | 25C/I/G/P |  |
|  | gm% | kcal% | gm% | kcal% | gm% | kcal% | gm% | kcal% |
| Protein | 18.8 | 20 | 19.0 | 20 | 19.3 | 20 | 19.3 | 20 |
| Carbohydrate | 61.1 | 65 | 61.6 | 65 | 62.7 | 65 | 62.7 | 65 |
| Fat | 6.5 | 15 | 6.5 | 15 | 6.7 | 15 | 6.7 | 15 |
| Total |  | 100 |  | 100 |  | 100 |  | 100 |
| kcal/gm | 3.78 |  | 3.81 |  | 3.88 |  | 3.88 |  |
| Ingredient | gm | kcal | gm | kcal | gm | kcal | gm | kcal |
| Casein | 200 | 800 | 200 | 800 | 200 | 800 | 200 | 800 |
| L-Cystine | 3 | 12 | 3 | 12 | 3 | 12 | 3 | 12 |
| Corn Starch | 390.5 | 1562 | 381 | 1524 | 362.3 | 1449.2 | 362.3 | 1449.2 |
| Maltodextrin 10 | 110 | 440 | 110 | 440 | 110 | 440 | 110 | 440 |
| Dextrose | 150 | 600 | 150 | 600 | 150 | 600 | 150 | 600 |
| Cellulose | 100 | 0 | 75 | 0 | 25 | 0 | 25 | 0 |
| Inulin * | 0 | 0 | 25 | 37.5 | 75 | 112.5 | 25 | 37.5 |
| Pectin ** | 0 | 0 | 0 | 0 | 0 | 0 | 25 | 37.5 |
| Glucomannan ** | 0 | 0 | 0 | 0 | 0 | 0 | 25 | 37.5 |
| Soybean Oil | 70 | 630 | 70 | 630 | 70 | 630 | 70 | 630 |
| Mineral Mix S10026 | 10 | 0 | 10 | 0 | 10 | 0 | 10 | 0 |
| Dicalcium Phosphate | 13 | 0 | 13 | 0 | 13 | 0 | 13 | 0 |
| Calcium Carbonate | 5.5 | 0 | 5.5 | 0 | 5.5 | 0 | 5.5 | 0 |
| Potassium Citrate, 1 H2O | 16.5 | 0 | 16.5 | 0 | 16.5 | 0 | 16.5 | 0 |
| Vitamin Mix V10001 | 10 | 40 | 10 | 40 | 10 | 40 | 10 | 40 |
| Choline Bitartrate | 2 | 0 | 2 | 0 | 2 | 0 | 2 | 0 |
| Yellow Dye #5, FD&C *** | 0.025 | 0 | 0.025 | 0 | 0 | 0 | 0 | 0 |
| Red Dye #40, FD&C | 0.025 | 0 | 0 | 0 | 0.025 | 0 | 0 | 0 |
| Blue Dye #1, FD&C | 0 | 0 | 0.025 | 0 | 0.025 | 0 | 0.05 | 0 |
| **Total** | **1080.55** | **4084** | **1071.05** | **4084** | **1052.35** | **4084** | **1052.35** | **4084** |
| * Inulin assigned caloric value of 1.5 kcal/gm (per Roberfroid, J. Nutr. 129: 1436S–1437S, 1999) |  |  |  |  |  |  |  |  |
| ** Pectin and Glucomannan assigned caloric value of 1.5 kca/gm (predicted) |  |  |  |  |  |  |  |  |

### Fecal and tissue sample collection

Fresh fecal pellets were obtained from mice on days 0, 7, and 35 and stored at -20 °C. On day 35, animals were anesthetized with isoflurane and then euthanized by cervical dislocation. Cecal and colon contents were collected, snap frozen in liquid nitrogen, and stored at -80 °C. Small intestine length, colon length, cecum weight, and gonadal fat mass were measured.

### DNA extraction

Genomic DNA was isolated from fecal pellets, cecal contents, and colon contents using a phenol-chloroform extraction method followed by a DNA clean-up. One fecal pellet, two colon pellets, or 100 mg of cecal contents were suspended in 750 μL of lysis buffer (200 mM NaCl, 100 mM Tris pH 8.0, 20 mM EDTA, and 20 mg/mL lysozyme) with 300 mg of 0.1 mm glass beads (BioSpec, Bartlesville, OK). Samples were vortexed briefly and incubated at 37 °C for 30 min. For each sample, 85 µL of 10% SDS and 20 µL of proteinase K (15 mg/mL) was added and the mixture was incubated for another 30 min at 60 °C. Bacterial cells were physically disrupted by adding 500 µL of 25:24:1 solution of phenol:chloroform:isoamyl alcohol and homogenizing in a bead beater (TeSeE Precess 48 Homogenizer, Bio-Rad, Hercules, CA) for 90 sec at 5300 rpm. Samples were then centrifuged at 13000 rpm for 5 min at room temperature. The resulting supernatant was further purified with sequential washes with 500 µL of 25:24:1 phenol:chloroform:isoamyl alcohol and 500 µL of 100% chloroform. Finally, DNA was precipitated with 100% ethanol and 3 M sodium acetate at -80°C for 60 minutes and then cleaned using the Qiagen QIAamp DNA Stool Mini Kit (Qiagen, Valencia, CA) per manufacturer’s instructions.

### 16S rRNA gene sequencing

Following DNA extraction, two consecutive polymerase chain reactions (PCR) were performed to amplify the V4 variable region of the 16S rRNA gene. The first PCR contained 120 ng of template DNA, six forward primers, six reverse primers (10 μM) and Robust DNA polymerase from the KAPA2G Robust PCR Kit (Kapa Biosystems, Wilmington, MA). Cycling conditions were: 95 °C for 3 min; [95 °C for 30 sec; 50 °C for 30 sec; 72 °C for 30 sec] x 10 cycles; 72 °C for 5 min. The second PCR further amplified the V4 variable region and added the Illumina MiSeq adapter primers and a single 12-nucleotide Golay error-correcting barcode for multiplexing (35). The KAPA HiFi HotStart ReadyMix reagent (Kapa Biosystems, Wilmington, MA) was used to amplify sequences from 5 µL of PCR product generated by the first reaction. Cycling conditions were: 95 °C for 3 min; [95 °C for 30 sec; 50 °C for 30 sec; 72 °C for 30 sec] x 22 cycles; 72 °C for 5 min. PCR products were purified using the HighPrep PCR clean-up kit (MagBio, Lausanne, Switzerland) and a DynaMag-96 side magnet (Life Technologies, Carlsbad, CA) per the manufacturer’s instructions. Final purified products were quantified and pooled at equimolar concentrations for high-throughput sequencing on the Illumina MiSeq platform (Illumina, San Diego, CA) at the University of North Carolina at Chapel Hill high-throughput sequencing facility.

### Characterization of the intestinal microbiota

16S rRNA gene sequence read classification was performed using the Quantitative Insights into Microbial Ecology 2 pipeline (QIIME2, version 2022.2) (36). Reads were demultiplexed and primers and adapters were removed. Divisive Amplicon Denoising Algorithm (DADA2) was used to correct for Illumina-sequenced amplicon errors and to generate absolute sequence variants (SV) at a 100% identity threshold (37). Sequences were truncated at 217 base pairs. To eliminate rare taxa, SV with lower than 0.01% of total reads were filtered out for each run. The number of SV decreased from 480 to 289 with 1,514,371 total reads after filtering for cecal samples. For colon samples, the number of SV decreased from 447 to 291, and the number of reads were reduced to 1,512,069. Fecal samples collected on day 35 of the study initially contained 580 SV of which 358 SV and 1,787,123 reads were retained after filtering. To compare across the three GI niches, unfiltered cecal contents, colon contents, and fecal samples at day 35 were merged and then filtered. The number of SV decreased from 790 to 391, with 3,892,532 reads after filtering. Ultimately, 29 cecal samples, 29 colon samples, and 30 fecal samples were successfully characterized and analyzed. The SILVA classifier (release 138, 99% OTUs) performed taxonomic assignments (38). MicrobiomeAnalyst (39) calculated α-diversity using Shannon diversity and the absolute number of SV, β-diversity using Bray-Curtis dissimilarity indices, relative abundance of bacterial genera sampled from the three GI niches, and differential abundance of bacterial genera detected in the cecum of mice on different diets with LEfSe (40). Multi-dimensional scaling (MDS) plots were generated in R (version 4.3.0) using the vegan (version 2.6.4) package.

### Statistical analysis

Average daily calorie intake, body weight change, gonadal fat mass, small intestinal and colon length, and cecum weight were compared between experimental groups using a one-way ANOVA with Tukey’s post-hoc test. PERMANOVA established statistical differences based on Bray-Curtis dissimilarity distances between diet groups and GI niches. A one-way ANOVA with Tukey’s post-hoc test assessed differences in α-diversity measures between diet groups and GI niche. Primary taxonomic analysis was carried out at the genus level (reflecting the most refined taxonomic level for these data). The top genera identified by LEfSe most likely to explain differences between groups and significant pairwise comparisons were determined by DESeq2 (version 3.18) (41).

---

**Supplemental Table 1.** LEfSe analysis of cecal microbial communities amongst diet groups.

**Supplemental Table 2.** FDRq values for pairwise comparisons between diets for the top 6 genera identified by LEfSe analysis.

